# Experience replay supports non-local learning

**DOI:** 10.1101/2020.10.20.343061

**Authors:** Yunzhe Liu, Marcelo G. Mattar, Timothy E J Behrens, Nathaniel D. Daw, Raymond J Dolan

## Abstract

To make effective decisions we need to consider the relationship between actions and outcomes. They are, however, often separated by time and space. The biological mechanism capable of spanning those gaps remains unknown. One promising, albeit hypothetical, mechanism involves neural replay of non-local experience. Using a novel task, that segregates direct from indirect learning, combined with magnetoencephalography (MEG), we tested the role of neural replay in non-local learning in humans. Following reward receipt, we found significant backward replay of non-local experience, with a 160 msec state-to-state time lag, and this replay facilitated learning of action values. This backward replay, combined with behavioural evidence of non-local learning, was more pronounced in experiences that were of greater benefit for future behavior, as predicted by theories of prioritization. These findings establish rationally targeted non-local replay as a neural mechanism for solving complex credit assignment problems during learning.

**One Sentence Summary:** Reverse sequential replay is found, for the first time, to support non-local reinforcement learning in humans and is prioritized according to utility.

## Main Text

Effective decision making incorporates new experience into our existing knowledge of the world. This allows us to determine the likely future consequences of different actions without having to experience it. When you encounter a traffic jam at a crossroads, for example, you learn that the route just taken should be avoided in the future, but you might also infer the value in avoiding the alternate paths that converge to this same location. Learning from direct experience can be straightforwardly achieved via “model-free” mechanisms that detect co-occurrence between actions (like routes taken), and subsequent rewards (*1*–*3*). However, it requires additional computation to propagate that experience to many other distal situations, as in the example of alternate converging roads. Despite behavioral evidence for this type of indirect learning, we understand little about how it is achieved in the brain (*4*–*7*).

In reinforcement learning (RL) theory (*8*), non-local value propagation can be achieved by “model-based” methods that leverage a learned map or model of the environment to simulate, or simply retrieve, potential trajectories (*9*, *10*). These covert trajectories can substitute for direct experience and thereby span the gaps between actions and outcomes (*11*), a process referred to as experience replay. A potential neural substrate for this process is the phenomenon of hippocampal “replay” in rodent. Here, cells in the hippocampus that encode distinct locations in space fire sequentially during rest in a time-compressed manner, recapitulating past or potential future trajectories (*12*–*14*). Utilizing methods developed to measure fast neural sequences noninvasively (*15*), such replay has now been found in humans during rest (*16*–*18*), with strong parallels to observations in rodents (*16*). Although these events appear appropriate to support value learning, there is little evidence in either species supporting their involvement.

If experience replay supports value learning, then its statistics should also be relevant for a second unresolved question: Given limited available time and resources, which of the myriad possible future actions does the brain prioritize during replay? A reward-maximizing agent should prioritize replay of whichever past experiences are most likely to improve future choices and thereby earn more reward (*19*). Recent theoretical analysis (*20*) argues that such rational priority for replay can be decomposed into the product of two factors, namely *need* and *gain*. *Need* captures how frequently a given experience will be encountered again in the future, while *gain* quantifies the expected reward increase from better decisions if that experience is replayed.

Accordingly, we designed a novel decision-making task that isolates the behavioral effect of nonlocal learning, while at the same time measuring neural signatures of replay and manipulating *need* and *gain*. We show that a specific subtype of neural replay is associated with non-local learning, and that replay of experiences with greater benefit for future behavior are prioritized, as predicted by RL theory.

### Task design

Our key hypothesis was that neural replay facilitates non-local learning, and that such replay is prioritized by its utility for future behavior. To detect human replay, we measured whole-brain activity using magnetoencephalography (MEG) while subjects performed a novel decision-making task. The task explicitly separates learning from direct vs. non-local experience, permitting the measurement of unambiguous neural and behavioral signatures of the latter (**Fig. 1A-C**).

**Fig. 1.**
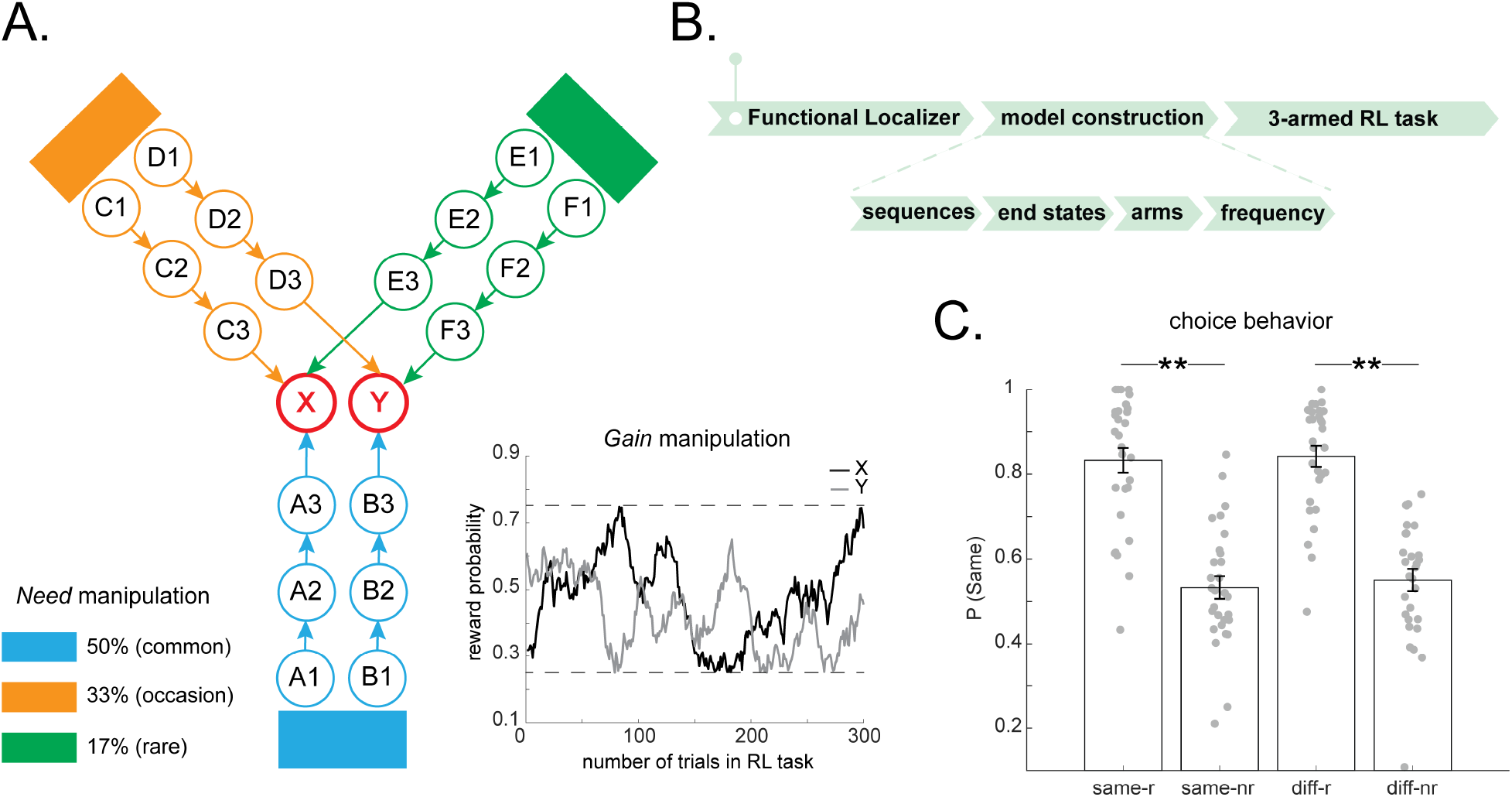
Experimental design for model-based reinforcement learning task. **(A)** The main RL task. At each trial, subjects were presented with one of the three starting states according to a fixed probability, and asked to select one from the two alternatives paths within this arm. The response associated with each alternative (left/right) was randomized across trials, and the reward probability of the outcome states varied slowly and independently over time. A crucial feature of this task is that outcomes (i.e., X and Y) are shared across all three arms, a design feature that enables non-local learning. *Need* is manipulated by the starting probability of each arm, shown on left. *Gain* is manipulated by the fluctuating reward probability of outcomes, X and Y respectively. An example of such reward schedule is shown on the right. The reward probability of X and Y changes gradually and independently over trials, with gaussian random walk, bounded between 25% and 75%. **(B)** Each experimental task phase is shown. Crucially, subjects learned the task model before commencing the main RL task. **(C)** Behavioral evidence showing exploitation of task model to aid learning. *Same/diff* is defined based on whether the current starting arm is the same or different to that of the last trial; *r/nr* indicates whether subjects were reward or not rewarded at the last trial. P(same) is the probability that subjects, in the current trial, select a path leading to the same outcome state as that in the last trial. Error bars shows the 95% standard error of the mean, each dot indicating results from each subject. * indicates *p* < 0.05, ** indicates *p* < 0.01.

To isolate local and non-local learning, the task comprised three starting states (henceforth called “arms”), each with two alternative choice paths (**Fig. 1A**). A choice at the starting state then leads to a sequence of three stimuli (“path”) followed by a final stimulus. At each trial, subjects are presented with one of the starting states and asked to make a choice between the two options with the goal of maximizing reward. Importantly, the two end states, reachable from each starting state, are shared across all three starting states. Each end state leads to a reward with a probability that changes slowly from trial to trial (**Fig. 1A**, **right)**. This task structure allows subjects to use reward feedback to inform their choices, including future choices at other starting states (non-local learning). This feature allows us to isolate learning about nonlocal options (since the local one may be driven at least in part by direct experience) as well as compare learning within-trial across nonlocal options with different properties (e.g., *gain* and *need*). The use of three-stimulus sequences allows unambiguous measurement of extended replayed sequences (vs co-occurrence) and their directionality.

In addition to distinguishing learning from local experience (the path just chosen) vs. non-local experience, the task allows us to test our hypotheses that replay and learning should favor the higher priority of the two non-local paths. Priority differed between paths as a function of both *need* and *gain*. Differences in *need* occurred because each starting arm was encountered with a different, but constant, probability: rare (17%), occasional (33%), and common (50%) respectively (**Fig. 1A**, **left)**. These probabilities were learnt prior to the main task (**Fig. 1B**). Since rewards were stochastic with fluctuating probability, the *gain* of propagating information about outcomes to different paths also fluctuated from trial to trial according to their individual reward histories. For instance, a newly encountered reward is more informative if this information promotes the selection of actions that would otherwise not be favored, whereas the absence of reward is more informative for avoiding actions that would otherwise have been chosen.

Thus, our task allowed us to investigate how subjects learn efficiently by incorporating new experiences, particularly those derived from a different starting state, into updated choices. To achieve this, subjects were first taught an overall task model comprising knowledge of the relations among different elements in the task, as well as the different starting probabilities assigned to each arm. To avoid any biased learning of the model, we introduced each component of the task carefully at different times (**Fig. 1B**).

### Functional localizer & Model construction

To index neural representations of states in the main RL task, we first showed subjects 18 visual stimuli in random order, a task phase called the *functional localizer*. These stimuli were later reused to form distinct states in the RL tasks (e.g., *A*1, *A*2, *A*3 in **Fig. 1A**). We constructed a probabilistic decoding model for each stimulus based on their evoked neural response in this *functional localizer* task. These decoding models are used later to search for sequential reactivation of states in the main RL task. Notably, these classifiers are unbiased to task experience and structure, because at the localizer phase subjects have no knowledge of the relationship among those stimuli, nor their value.

The experiment proceeded across distinct phases to ensure knowledge of the task model (i.e., *model construction*, **Fig. 1B**). Upon completion of a functional localizer, subjects learned how the 18 stimuli formed 6 distinct sequences, i.e., the relationship among the 18 stimuli. We refer to this as *sequence learning.* Subjects next learned a mapping between sequences and end states, i.e., *end state learning* and subsequently learned which sequence belongs to which starting arm, i.e., *arm learning*. Up to this point, experience is still unbiased as subjects have only learnt the relational structure between arms, end states, and sequences (i.e. no reinforcement). At the end of this stage, subjects learned the starting probability of each arm, including the fact that these probabilities remain constant throughout the experiment. Subjects also learned the frequency of each starting arm by experience, i.e., *frequency learning*. To ensure subjects had acquired knowledge of the full task structure we included quiz after each learning phase. Subjects’ performance was always above 85% (see details of training and task in Materials and Methods). Upon completion of the entire model construction stage, subjects performed the main RL task (**Fig. 1A**).

### Behavioral evidence of non-local learning and prioritization

In the RL task, subjects need to learn the value of each action at each starting arm, with the aim of maximizing reward. Direct, model-free learning allows subjects to favor a previously rewarded action when they encounter the same starting arm. Accordingly, when the starting arm in one trial matched that of the previous trial, we found that subjects were more likely to repeat the same action if they had been rewarded compared to not rewarded on the last trial (Mixed effects logistic regression, *p* = 7.5×10^−15^). We then tested whether subjects transfer the value obtained in the chosen (i.e., local) path to the other non-local paths that lead to the same outcome. Achieving effective non-local learning requires use of a model-based mechanism, such as replay, to propagate local rewards to non-local actions. We found that a path leading to a previously rewarded end state was favored even when the choice was presented at a different starting arm (*p* = 9.5×10^−23^), and this effect did not differ significantly between trials whether the starting arm was repeated or not (*p* = 0.90 for the main effect of arm, *p* = 0.46 for the interaction effect between arm and reward, **Fig. 1C**). This is a hallmark of non-local, model-based learning (*4*, *21*).

The previous analyses consider choices only as a function of events on the single preceding trial. To ask more detailed questions about learning, we built a computational model that also incorporates longer-run effects of experience on multiple later choices. In particular, we fit trial-by-trial behavior to a modified Q-learning model (*22*), that updates the value of each action from obtained rewards, and chooses actions on the basis of this value (see Materials and Methods for modelling details). By separating learning for local (*α*_*d*_), vs. non-local paths (*α*_*n*_), we found non-local action values were updated to a similar extent as local action values (*α*_*d*_ = 0.64; *α*_*n*_ = 0.60; difference in learning rates = 0.04, *p* = 0.61), consistent with the model-agnostic results.

To ask whether the behavioral signature of learning from non-local outcomes was greater for paths with higher priority, we augmented the baseline model with additional free parameters measuring the strength of non-local learning as a function of each of the two features that determines priority: *gain* (the informativeness of the current reward for improving choice at the replayed arm) and *need* (the likelihood that arm will be visited in the future, given by its frequency). This was possible because, in the task, there are always two non-local paths sharing the same end state with the current chosen one, allowing us to compare learning directly across them. We calculated the strength of learning by estimating separate learning rates for the higher and lower priority paths on each trial, in addition to a third learning rate for updating the local (chosen) path (*α*_*d*_ = 0.63). Numerically, a higher learning rate was estimated for both higher-gain (*α*_*h*_= 0.79 vs *α*_*l*_= 0.37, **Table. S1**) and higher-need paths (*α*_*h*_= 0.61 vs *α*_*l*_= 0.54, **Table. S1**), a difference significant for gain (*p* = 0.020) but not need (*p* = 0.16, **Table. S1**). These results provide behavioral support for a hypothesized rational prioritization of non-local learning.

### Neural decoding & Sequential reactivations

We turned next to neural data to ask how the observed non-local learning is achieved in the brain. First, we verified we could decode all 18 visual stimuli well above chance, with a peak cross-validation decoding accuracy at 47 ± 3 % (vs. chance level, 1/18 ≈ 6%), based on evoked neural response in the *functional localizer* task (**Fig. 2A, B**, see also **Fig S1**, and Materials and Methods for decoding analysis details). By applying the decoding models of these 18 stimuli to the RL task, we could test for their sequential reactivations at the point of outcome receipt, the time period when new learning can occur. Note, the focus on this period is analogous to the time when rodents consume a reward and backward replay sequences are often observed (*23*) (but also see discussion for connections to rodent sequences). We operationally refer to any reactivation of sequences here as replay, given we are looking for patterns of off-task spontaneous reactivations.

**Fig. 2.**
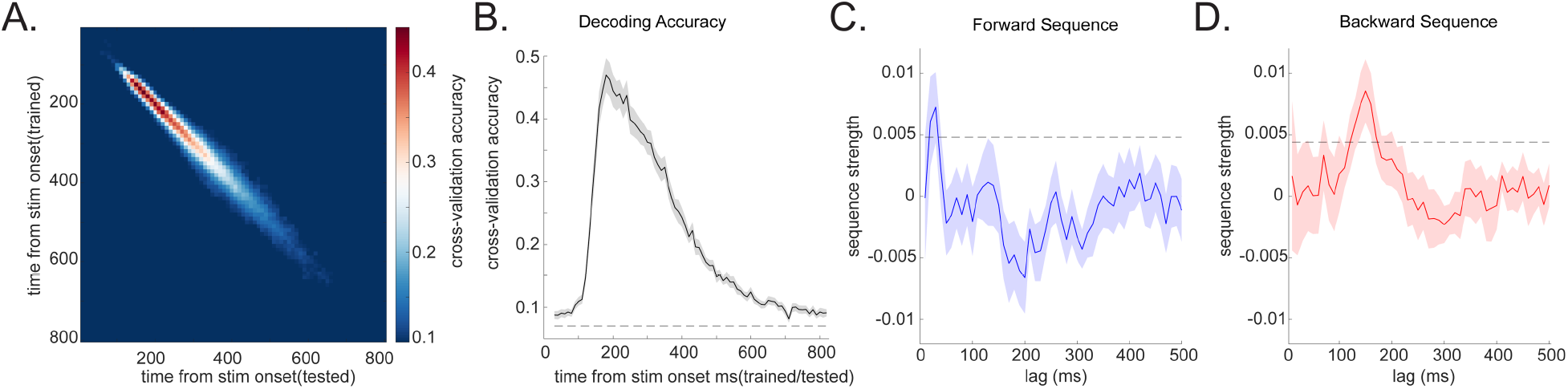
Stimuli decoding and neural sequences. **(A)** All 18 stimuli classifiers are trained based on their evoked multivariate neural patterns at each time bin, from 10 ms to 800 ms post stimulus onset, in the *functional localizer* phase and tested at all time points across 10-800 ms in a leave-one-out cross validation scheme. This gives us temporal generalization plots, with the Y axis indicating the time bins classifiers were trained on, and X axis indicating the test time of classifiers. Accuracy was obtained from all 18 stimuli classifiers, and the readout is deemed accurate if the corresponding classifier of the test label gives the highest decoding probability, as in previous studies (*16*, *17*, *24*). The diagonal of the temporal generalization plots is the decoding accuracy at the same time we trained the classifiers on, peaking at approximately 200 ms post stimulus onset. **(B)** We trained stimuli classifiers based on their evoked neural response at 200 ms, as in previous studies (*16*, *17*). The dotted line is the permutation threshold taken as the 95% percentile of peak decoding accuracy on randomly permuted labels. (**C-D**) Applying trained classifiers to time of outcome receipt in the RL task. A sequence analysis (*15*) provided evidence for two distinct sequence signatures, one forward sequence (blue) peaking at a 30 ms state-to-state time lag (C), and a backward sequence (red) peaking at 160 ms time lag (D). The dotted line is the permutation threshold that controls for multiple comparisons. It was taken as the 95% percentile on the peak sequenceness value over all computed time lags in permutation. This permutation is done by randomly permutating the transition matrix, which are shown to be statistically robust (*15*–*17*). The X axis is the time lags. Sequence analysis is done separately at each time lag. The Y axis is the evidence of sequenceness, i.e., sequence strength.

We first look for spontaneous replay of all possible transitions consistent with the model. We refer to sequences that express the same direction as experience (e.g., *A*1 → *A*2 → *A*3) as forward replay, and sequences in the opposite direction (e.g., *A*3 → *A*2 → *A*1) as backward replay. Utilizing a recent methodological advance in MEG decoding of replay, we assessed evidence for replay in a forward and backward direction separately and at different speeds (*15*). We found significant forward replay peaking at 30 ms state-to-state time lag (**Fig. 2C**), and backward replay that peaked at 160 ms state-to-state lag (**Fig. 2D**, see Materials and Methods for sequence analysis details). Consequently, we focus on this 30 ms forward and 160 ms backward replay in all later analyses.

### Two types of replay: functional and physiological differences

Our observation of forward replay with 30 ms state-to-state time accords with previous work measuring replay in humans during post-task rest (*16*), but extends on those findings to a context that now includes learning. The 160 ms backward replay has not been reported previously, but it is intriguing as its direction is consistent with proposals for solving credit assignment by backpropagating reward, based on theory (*20*) and empirical data (*12*, *16*, *23*). If this 160 ms backward replay supports non-local updating, we would expect it to also represent the contents of non-local paths. In line with this prediction, we found that the 160 ms backward replay dominantly represents non-local paths (one sample *t* test, *t* (28) = 2.92, *p* = 0.007), and to a significantly greater degree than local ones (paired *t* test, *t* (28) = 2.21, *p* = 0.03, **Fig. 3A**). The 30 ms forward replay shows an opposite pattern (interaction between replay types and representational content, *F* (1,28) = 15.01, *p* = 0.001), not significantly representing the non-local paths (one sample *t* test, *t* (28) = −0.09, *p* = 0.93), but mainly the local one, corresponding to recent experience (*t* (28) = 3.37, *p* = 0.002, **Fig. 3B**).

**Fig. 3.**
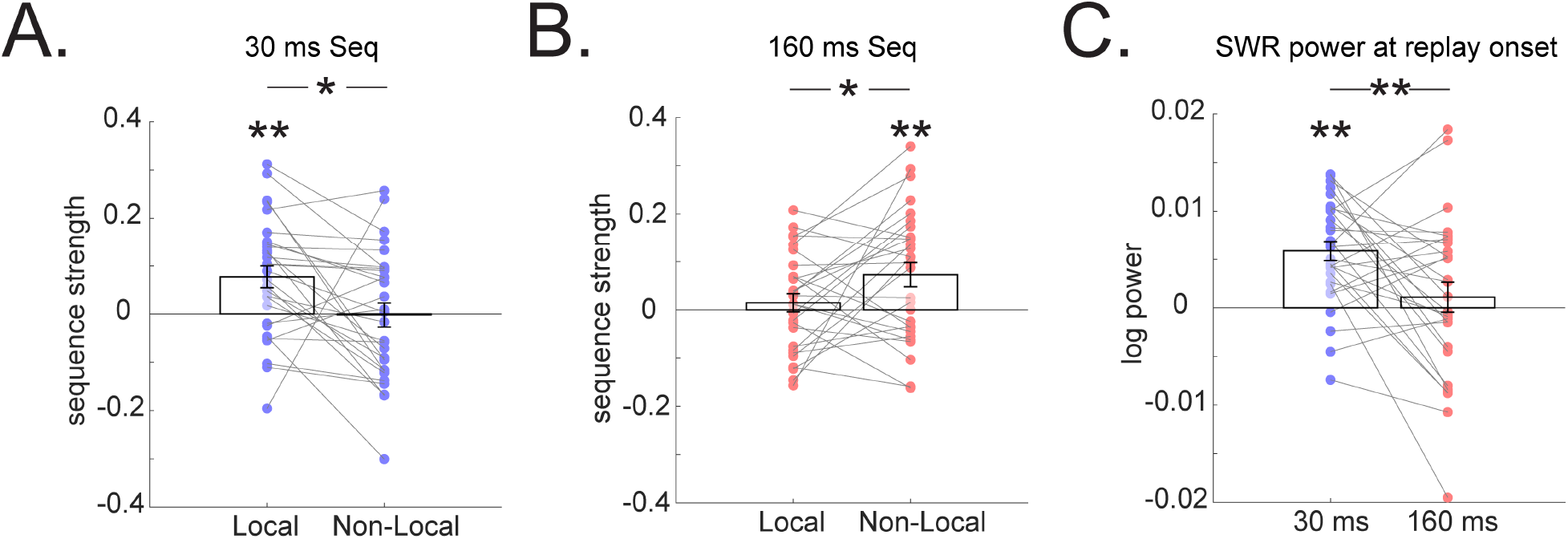
Representational and physiological differences between the two types of replay. **(A)** A 160 ms backward replay encodes non-local as opposed to local experience. **(B)** A 30 ms forward sequence encodes local experience alone, but not non-local. **(C)** The initialization of 30 ms forward sequence is associated with a power increase in a ripple frequency band (120-180 Hz), consistent with previous study (*16*), but this is not the case for 160 ms backward sequence. These frequency power signatures are significantly different. The grey line connects results from the same subject. Error bars shows the 95% standard error of the mean, each dot indicating results from each subject. * indicates *p* < 0.05, ** indicates *p* < 0.01.

In addition to a representational difference between the two types of replay, we also tested whether these distinct replay signatures differ in terms of their underlying physiological properties. Our previous study (*16*) showed that fast human replay (with time lag of 20-50 ms) during rest is associated with an increased ripple frequency (120 −150 Hz) power, akin to sharp wave ripple replay in rodents (*25*–*27*). We replicated this finding here, showing that initialization of a 30 ms forward replay is associated with a ripple frequency power increase (one sample *t* test, *t* (28) =5.82, *p* = 2.9×10^−6^), but this power increase is not seen for the 160 ms backward replay (*t* (28) =0.71, *p* = 0.48). A significant difference was evident in the power of ripple frequency between the two types of replay (paired *t* test, *t* (28) =2.84, *p* = 0.008, **Fig. 3C**). Beamforming results indicated that while both replay types are associated with activation in visual cortex and medial temporal lobe, a 30 ms forward replay has higher hippocampal activation compared to the 160 ms backward replay, while conversely, the 160 ms backward replay has greater cortical engagement (**Fig S2**).

### Non-local Replay facilitates non-local learning

Having identified neural replay candidates for learning, we then tested whether non-local replay (i.e., the 160 ms backward replay) facilitates non-local learning in choice behavior and, if so, whether such replay is competitively prioritized in accord with theoretical accounts (*20*). We again posed these questions in terms of RL-based computational models of trial-by-trial choice behavior (see Materials and Methods for modelling details).

First, to ask whether replay helps non-local learning, we augmented the baseline Q-learning model with a term measuring learning in the presence vs absence of trial-by-trial neural replay. In particular, having first separated learning rates for local and non-local paths (as before, these are paths leading to the same end state), we tested whether the baseline learning rate for each non-local path was significantly increased on trials when that path expressed significant neural replay, vs. not. We found higher nonlocal learning rates in the presence vs. absence of significant 160 ms backward replay (see supplementary material for details) (*α*_*replay*_ = 0.70; *α*_*no-replay*_ = 0.61; difference in learning rates = 0.09, *p* = 0.023, **Table S2**). This was not the case when we repeated the same analysis for the 30 ms forward replay (difference in learning rates = 0.01, *p* = 0.457, **Table. S2**), as expected given the lack of representation of non-local paths in the 30 ms forward replay.

Finally, we asked whether replay is prioritized to favor more useful experiences. Consequently, on each trial, we compared the strength of replay across the higher vs. lower priority non-local paths, using a net priority score incorporating the product of *gain* (estimated per-arm, -trial, and -subject from the behavioral model) and need (17%, 33%, 50%, for paths in rare, occasion and common arm, respectively) (**Fig. 4A**). We observed significantly greater replay for the higher priority path, evident for backward replay at 160 ms time lag (*t* (28) = 3.30, *p* = 0.003), and with no effect for 30 ms forward replay, *t* (28) = −0.34, *p* = 0.74, **Fig. 4B**). All these results are specific to paths leading to the same end state (where non-local learning happens). No facilitation or prioritization of either type of replay was seen for paths leading to a different end state (**Fig. S3**). Model-agnostic analyses (based on comparing replay as a function of reward and need) parallel these results (**Fig. S3**).

**Fig. 4.**
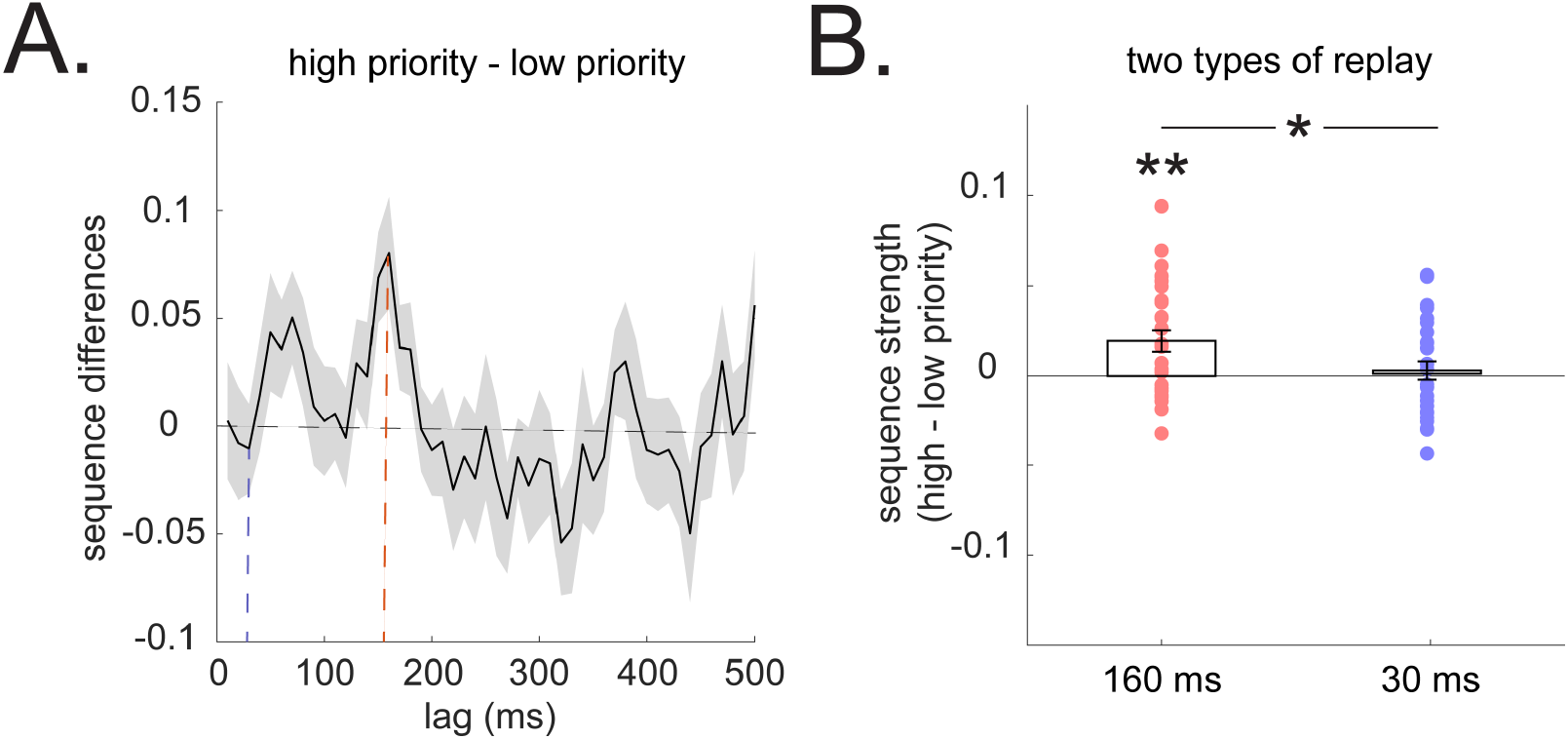
Prioritization of non-local replay. **(A)** Sequenceness differences in high vs. low priority (determined by *gain*×*need*) of non-local paths. We use a backward minus forward sequenceness to provide a summary replay statistic at each time lag. A positive value indicates greater backward sequenceness, while a negative value indicates higher forward sequenceness, similar to previous studies (*16*, *17*). This is for visualization only. The above-zero peak at 160 ms time lag suggests a greater backward replay for a high priority path. The dotted blue line denotes result at 30 ms lag, and the dotted red line indicates result at 160 ms lag. **(B)** The 160ms backward sequence alone is replayed more in a higher priority non-local path compared to lower priority one. The 30 ms forward replay does not differentiate the two non-local paths. Error bars shows the 95% standard error of the mean, with dots indicating results from each subject. * indicates *p* < 0.05, ** indicates *p* < 0.01.

## Discussion

In current study, we disassociate between two types of replay as a function of local vs. non-local learning, establishing for the first time, a connection between neural replay and learning via a non-local credit assignment as expressed in behavior.

We found that replay facilitates learning of action values by recapitulating non-local experiences, evidenced as enhanced learning effects on subsequent choices. In other words, replay connects actions and outcomes across intervening states, offering a neural mechanism for model-based RL. Furthermore, we demonstrate that the content of this replay, and separately the strength of updating as expressed behaviorally, are prioritized according to their utility for future behavior, as predicted in RL theory (*20*).

These findings corroborate a long-standing hypothesis about the role of awake replay on model-based planning and credit assignment, an hypothesis heretofore based primarily on rodent studies reporting replay patterns that would be appropriate for this function, but without linking directly to behavioral changes (*12*, *23*). These results also extend on our previous fMRI results in humans linking non-local reactivation (without assessing sequences) to planning (*4*, *5*, *28*). In the current study, by exploiting the temporal resolution of MEG and the use of three-stimulus sequences, we could distinguish sequential replay from reactivation of isolated states. Notably, we observed no significant effects related solely to reactivation of individual states (**Fig. S4**).

The 160 ms backward replay supporting non-local learning is distinct from the forward 30 ms replay reported in previous studies (*16*, *24*). Unlike the latter, the 160 ms replay is not associated with a ripple frequency power increase (*16*). This raises an intriguing possibility that the 160 ms replay, which has a state-to-state transition frequency of around 6 Hz, might be a homologue of rodent theta sequences or similar (*29*–*32*). However, theta sequences generally occur during ongoing behavior in rodents and are in a forward direction, akin to a “look ahead” signal (but see (*33*) for backward theta sequence), while the 160 ms sequence we identify is backward in direction and occur at a trial’s end.

Whether or not the reverse replay corresponds to either rodent phenomenon, its backward direction, representational contents and timing are suited to solve the non-local credit assignment problem, where an outcome at the end of a path impacts on decisions made at the (alternative) beginning. Theoretical work has focused more often on forward replay (or mental stimulation) of potential trajectories assumed to occur at choice time. Such patterns – more reminiscent of “planning” in the colloquial sense – also occur in rodents and could also, in principle, solve the current task. More generally, in the same framework they can be viewed as another means by which replay serves to connect actions and outcomes (*20*). In the current study, we found no evidence that forward replay at choice time related to credit assignment (**Fig. S5**). Such a process may play a role in other circumstances or in other task implementations, for instance in games like chess where particular choice situations are unlikely to have been anticipated ahead of time.

Our findings reveal that non-local experience replay supports planning and, more broadly, it serves as a neural mechanism crucially involved in ‘model-based reinforcement learning’. Accordingly, our findings extend the role of replay to account for non-experiential inferential learning and are remarkably consistent with reinforcement learning theory.

## ACKNOWLEDGMENTS

We thank Rani Moran and Evan Russek for discussions on study design and analysis.

## Funding

This work was supported by a Wellcome Trust Investigator Award (098362/Z/12/Z) to R.J.D. This work was carried out whilst R.J.D. was in receipt of a Lundbeck Visiting Professorship (R290-2018-2804) to the Danish Research Centre for Magnetic Resonance. Y.L. is supported by a UCL Graduate Research Scholarship and a UCL Overseas Research Scholarship. Y.L. is a pre-doctoral fellow of the International Max Planck Research School on Computational Methods in Psychiatry and Ageing Research (IMPRS COMP2PSYCH). T.B is supported by a Wellcome Trust Senior Research Fellowship (104765/Z/14/Z), and a Principal Research Fellowship (219525/Z/19/Z), together with a James S. McDonnell Foundation Award (JSMF220020372). N.D and M.M are funded by the US National Science Foundation grant IIS-1822571, part of the CRCNS program.

## Author Contributions

Y.L, M.M, T.B, N.D & R.J.D contributed to conception and design of the study; Y.L contributed to data acquisition with help from M.M; Y.L, M.M & N.D contributed to data analysis. Y.L wrote the manuscript with assistance from all authors.

## Competing interests

None.

## Data and materials availability

Data used to generate the findings of this study will be freely available upon request (subject to participant consent) to the Lead Contact. Full analysis code will be made available upon publication. Simulation code for sequenceness analysis can be found in https://github.com/YunzheLiu/TDLM

## Materials and Methods

### Participants

31 adults aged 19–31 participated in the experiment. They were recruited from the UCL Institute of Cognitive Neuroscience subject pool and from a mailing list for UCL MSc students. Eighteen were female, and three left-handed. All participants had normal or corrected-to-normal vision and had no history of psychiatric or neurological disorders. Two subjects were excluded from later analysis - one was a pilot subject and had gone through a slightly different task procedure of task, the other subject had metallic material in the hair, making the MEG data unusable. Thus, 29 subjects contributed data for the reported results. All participants provided written informed consent and consent to publish prior to start of the experiment, which was approved by the Research Ethics Committee at University College London (UK), under ethics number 1825/005.

### Overview of the task design

The task comprised three phases: functional localizer, model construction, and a 3-armed reinforcement learning (RL) task. The task and training regime were designed specifically to avoid bias in state decoding and ensure participants learned a correct model of the relational structure before the RL task. The task was implemented in MATLAB (MathWorks) using Cogent (Wellcome Trust Centre for Neuroimaging, University College London).

### Functional localizer task

The 18 distinct visual stimuli were shown in a randomized order. Those stimuli corresponded to intermediate states in the main RL task, forming 6 distinct sequences with no overlapping state representation (e.g., *A*1 → *A*2 → *A*3; *B*1 → *B*2 → *B*3, etc). Note that the mapping between stimuli and states was randomized across subjects. We trained one classifier for each stimulus based on its evoked neural response in the localizer task and used the resulting set of classifiers to decode their reactivations in the later RL task.

At each localizer trial, a visual stimulus (e.g., house) was presented at the centre of the screen for 800 ms, and participants were asked to think about its semantic meaning, i.e., this is a house. After that, a text label (e.g., “house” or “face”) appeared, and the subjects were asked to make a yes or no response within 1000 ms, using a response box (mean accuracy: 97.3 ± 0.4%, Mean ± SE). This was followed by a jittered 500-700 ms inter-trial interval (ITI). There are 50 trials for each stimulus presentation, resulting in 900 trials in total. This task is designed to encourage semantic representation of the stimuli, which we have found useful for sequence detection in previous studies (*16*, *17*).

### Model construction task

As we are interested in studying model-based or nonlocal learning we first need to ensure participants learnt the correct task structure (i.e. a “model” or map of the non-local relationships among stimuli in the task). Subjects were taught task structure in the following order: 1) *sequence learning*: stimuli-sequence mapping, 2) *end state learning*: sequence-end state mappings; 3) *arm learning*: sequence-arms mapping, 4) *frequency learning*: frequency of occupying each of the three different task arms.

During *sequence learning*, participants learnt how 18 visual stimuli form 6 distinct sequences, where each sequence comprised 3 stimuli. The 3 stimuli appeared on the screen in the correct task order (e.g., *A*1 → *A*2 → *A*3), with each stimulus in a sequence presented for 1000 ms. There were 3 learning blocks with each sequence presented 10 times within each block. To test whether participants learnt the within sequence transitions, we probed their knowledge at the end of each block. At probe trials, all three stimuli from the same path were presented, but in a scrambled order. Subjects were required to select these stimuli in true order sequence. Following learning, mean *within* sequence knowledge accuracy was 98.0 ± 0.8%. We also tested *between* sequence knowledge, by presenting one stimulus alone on screen for 1000 ms and asking participants to reflect on which path it belonged to. After that, two alternative stimuli were presented: one drawn from the same path as the probe stimulus, another drawn from a different path. Subjects were required to choose the stimulus that belonged to the same path. The mean accuracy in this probe task was 92.1 ± 0.9%. Together, these results suggest participants learnt the mapping between stimuli and sequences.

At *end state learning*, participants learnt which sequence leads to which end state (*X* or *Y*). Here it was crucial that subjects understood each end state was shared across three distinct paths, such that half of all paths led to the same end state, *X*, while the other half led to the alternative end, *Y*. During learning, stimuli in each sequence were shown sequentially, followed by the end state. Each sequence and end state mapping were repeated 10 times. After that, we tested subjects’ knowledge. In a probe trial, one sequence (consisting of three stimuli) was shown on screen, and subjects were required to think about which end state it led to for a total of 1000 ms. Two end states were then shown on the screen, and subjects were required to select the correct one within 1000 ms. The mean accuracy was 97.1 ± 0.8% in mapping from sequence to end. We also tested a mapping the other way around, i.e., from end state to sequence. Here the end state was shown at centre screen for 1000 ms and subjects were required to think about which sequences led to this end. After that, two sequences were shown on the screen where these were associated with different end points, and participants were asked to select the one that lead to the same end within 1000 ms. To avoid participants relying on a single stimulus alone (because each sequence comprises three distinct stimuli) when making a choice, one out of three stimuli in a sequence was randomly blocked off visually at each probe trial, and this encouraged participants to think about the sequence as a whole. Mean accuracy was 94.1 ± 0.9%. Overall, these results suggest participants learnt a mapping between sequences and end state.

In the *arm learning* phase, participants acquired knowledge of which sequences belong to which arms. There were three starting arms, and each arm had two sequences. The three starting arms had different frequencies in the main RL task, but in this phase all arms were experienced equally. Importantly, participants also did not know yet which arm they would encounter often or rarely in the main RL task, as this information was only learnt in the next phase. In this phase, the learning procedures were similar to the *end state learning* phase. After learning, the participants showed good knowledge regarding both the mapping from sequence to starting arms (mean accuracy is 95.4 ± 1.0%), and from starting arms to sequences (mean accuracy is 92.2 ± 1.0%), suggesting they had also learnt a mapping between sequences and starting arms.

In the *frequency learning* phase, participants learnt the encounter frequency associated with each starting arm. This was fixed and not chosen by the subjects. The mapping between arms and frequency was randomized across subjects but fixed within subjects. It indicated how likely each trial might start in any given arm. These different starting probabilities aimed to create different *need* level in the RL task, and is akin to successor representation in reinforcement learning literature (*34*). We informed participants explicitly the starting probabilities of each arm, i.e., rare - 17%, occasion - 33%, common - 50%. Additionally, we let participants experience the probability differences by showing the three arms according to their encountering frequency in the later RL task. After that, participants were quizzed on the mapping between arms and their starting probabilities. The mean accuracy is 95.9 ± 2%, suggesting they learnt the frequency differences among the three starting arms.

### Three-armed RL task

After subjects acquired all the necessary knowledge about task structure, they then performed the main RL task. This consisted 5 blocks, each block containing 60 trials, resulting in 300 trials in total. At each trial, a starting arm was shown first, chosen pseudo-randomly according to the starting probabilities. The arm picture appeared for 2500 ms, during which the participants were required to think about which one out of two paths within this arm they wished to choose. After that, the first stimulus of the two paths was shown on screen (e.g., *A*1 & B1, with their left/right location randomized across trials). Participants were allowed up to 1000 ms to enter a response indicating their choice. The chosen path was then played out, with each stimulus appearing sequentially at centre screen for 500 ms. This sequence presentation was followed by the end state, at which point participants were asked to press the “advance” key to reveal its value (£1 or 0). This disassociated visual offset of the end state picture from the reward period. The value display lasted for 2500 ms and was followed by a jittered 500-700 ms ITI. The output of the two outcome states was binary and independent from each other. The reward probability for each outcome state followed an independent Gaussian random walk with mean equal to zero and standard deviation equal to 0.2, bounded between 0.25 and 0.75 with reflecting boundaries, similar to that used in previous studies (*21*, *35*).

At the end of each block, we tested the subject’s knowledge of the transitions within sequences in order to ensure they had not forgotten those transitions along the experiment. Participants were again shown three stimuli from the same sequence, but in a scrambled order, and were asked to select the stimuli in the right sequence order. They were tasked to do this twice for each sequence. The mean accuracy is 96.2 ± 0.5%, and this did not change as a function of blocks (*F* (4,112) = 0.89, *p* = 0.46), suggesting sequence knowledge was preserved across the whole RL task.

### MEG Acquisition and Pre-processing

We recorded whole brain neural activity using magnetoencephalography (MEG) throughout the experiment, except for the time participants were experiencing the frequency of different starting arms prior to the RL task. MEG was recorded continuously at 1200 samples/second using a whole-head 275-channel axial gradiometer system (CTF Omega, VSM MedTech), while participants sat upright (3 sensors not recorded due to excessive noise in routine testing). The task was projected onto a screen suspended in front of participants, and participants made responses on three buttons of a MEG-compatible button box, indicating “up/left,” “down/right,” and “advance” respectively.

Preprocessing was conducted separately for each scanning session, as in our previous study (*16*). Sensor data were high-pass filtered at 0.5 Hz using a first order IIR filter to remove slow-drifts. Data were then resampled to 100 Hz (decoding, reactivation and sequenceness analysis) and 400 Hz (time-frequency analysis), and excessively noisy segments and sensors removed before independent component analysis (ICA). ICA (FastICA, http://research.ics.aalto.fi/ica/fastica) was used to decompose the sensor data for each session into 150 temporally independent components and associated sensor topographies. Artefact components (e.g. eye blink and mains interference) were classified by automated inspection of the spatial topography, time course, kurtosis of the time course and frequency spectrum for all components. Artefacts were rejected by subtracting them out of the data. All analyses were performed on the filtered, cleaned MEG signal at whole-brain sensor level.

### Behavior Analysis

To test the behavioural effects of local and nonlocal learning from rewards in a relatively model-agnostic fashion, we used a mixed-effects logistic regression. In particular, we analysed the choice made on each trial (coded as stay vs switch, with respect to whether the choice leads to the same end state, e.g. X, as the path chosen on the previous trial). The chance of staying was analysed as a function of two binary factors: whether the previous trial was rewarded or not, and whether the current trial started in the same, or a different, start state as the previous trial.

The logic is that a reward (vs non-reward) should increase the chance of staying with the choice of the same end state on the next trial; but this effect can be learned by direct experience only if the start state for the new choice is the same as that from which the reward was received. Thus, if participants behaved in a purely model-free way, they would not be able to use the outcome in the last trial to inform a decision in the current trial if it now involves a different starting arm. To favour the path leading to an option that was rewarded when starting from a different starting state consistent with such a model-based learning account. This logic parallels a two-step decision-making task that has been used to measure model-based vs. model-free contributions to choice (*4*, *21*). We have also simulated choice behavior that recapitulated the same reward schedule and starting arms probabilities as in the RL task using a model-free vs. model-based Q learning model (*4*, *21*). Simulation results support the line of reasoning we have outlined above.

We estimated the model using generalized linear mixed-effects logistic regression, including as explanatory variables the two factors and their interaction. All four coefficients of the model (the intercept, two main effects, and interaction) were all taken as random effects (allowed to vary across participants).

### Behavior Modelling

To provide a more-detailed model of learning on the task, which examines how each choice is driven by rewards received potentially across multiple preceding trials, we built a series of computational RL models (*20*). These are all based on a baseline model that combines the possibility of both local (model-free) and non-local (model-based) learning steps, nesting both simpler models as special cases. In particular, the model assumes that participants’ choices are guided by a value function *Q*(*s, a*) estimating the chance of reward for each of the two actions *a* available from each of the three starting states *s*. The choice on each trial is modelled as softmax in these estimates, i.e. *P*(a_t_ = *a*|*s*_*t*_ = *s*) ∝ *βQ*(*s*, *a*), with free inverse temperature parameter *β*.

Following each trial, the estimates *Q* are then updated according to the reward received. Importantly, on each task trial there is one directly experienced path as well as two non-local paths that lead to the same end state. We allow for both local learning (about the value *Q*(*s*_*d*_, *a*_*c*_) of the action chosen *a*_*c*_ on the current trial at the actual start state *s*_*d*_) and nonlocal updates (about the value of the equivalent action at each of the other two start states, *s*_*n*1_ and *s*_*n*2_ (Here the action is defined in terms of the end state it leads to, so that, if one takes the same action, i.e., *a*_*c*_ in the nonlocal arms, the outcome is the same end state.). This leads to three steps of learning on each trial

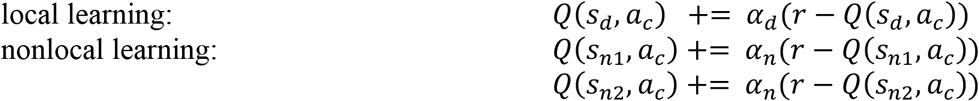

The updating happens only in paths that lead to the same end state, because the reward schedule for each outcome is independent – i.e. getting reward (or not) at *X*, does not provide any information about *Y*. Importantly, this baseline model includes separate learning rates, *α*_*d*_ and *α*_*n*_, controlling the speed of direct vs. non-local learning respectively. By estimating this model, we can measure the strength of nonlocal learning separately, where *α*_*n*_ = 0 recovers a simpler direct learning model.

The remaining behavioural analyses are all based on further refinements of this basic model, which separate the two nonlocal starting arms on each trial, *s*_*n*1_ and *s*_*n*2_, according to different properties that might affect learning, estimate separate learning rate parameters for each, and compare them. Thus, we first tested whether learning was faster for the higher (*s*_*h*_) vs lower (*s*_*l*_) priority arm of the two nonlocal arms on each trial. This replaces the previous parameter *α*_*n*_ with two learning rates, *α*_*h*_ and *α*_*l*_. The updating rules are then:

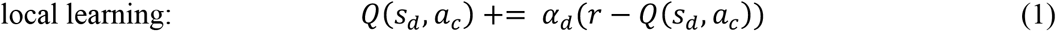

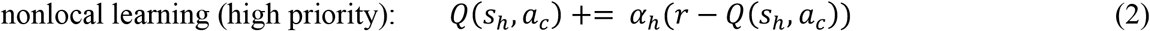

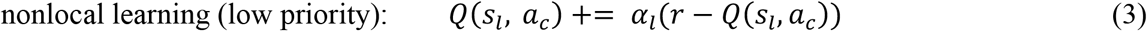

We test two versions of this model, where high and low priority are defined in terms of either need or gain. For prioritization based on *need*, this calculation is straightforward because the starting probability for each arm is fixed and known to the participants. Consequently, *s*_*h*_ has simply defined as the arm with the higher starting probability of the pair. Prioritization based in *gain* is more complicated, because gain (formally, how much reward will be improved if an update is performed and the state is revisited) depends on the currently estimated *Q* values at each state. We incorporate a trial-by-trial measure of gain into the model and use it to assign *s*_*h*_ and *s*_*l*_. Based on Mattar and Daw (*20*) the gain can be written:

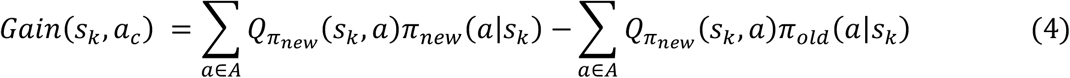

Where *s*_*k*_ indicates a non-local arm. *π*_*new*_(*a*|*s*_**k**_) represents the probability of selecting action *a* in state *s*_k_ after a learning step, and π_*old*_(*a*|*s*_*k*_) is the same quantity prior to the learning step. In this task, there are only two actions available in *A*: *a*_*c*_ and ¬*a*_*c*_, where ¬*a*_*c*_ indicates the action that leads to the alternative end state.

Assuming that if a learning step occurs, it is full, and a softmax decision policy, the above Equation(4) can be re-written based on reward information in current trial:

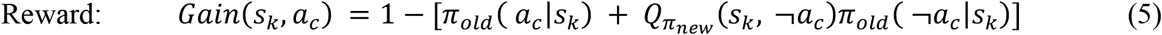

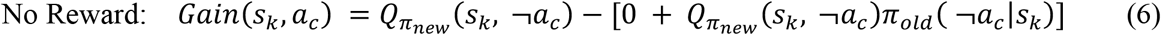

Where 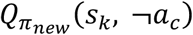 is the same as 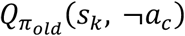 given no updating happens at the action leading to a different end state. We can therefore compare *Gain* in two non-local arms and assign *α*_*h*_ to a higher gain path, and *α*_*l*_ to lower gain path. In the end, we are interested whether *α*_*h*_ > *α*_*l*_.

To estimate the models, we utilized Markov Chain Monte Carlo (MCMC) methods, implemented in the Stan modelling language (Stan Development Team). We produced 4 chains of 10,000 samples each. The first 2500 samples from each chain were discarded to allow for equilibration. We verified the convergence of the chains by visual inspection, and additionally by computing for each parameter the ‘potential scale reduction factor’, 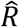 (*36*). For all parameters, we verified that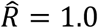, consistent with convergence (*37*). As a sanity check, we also ran parameter recover to ensure the modelling results are not biased due to certain reward schedule.

The model was specified hierarchically, so that participant-specific parameter estimates were assumed to be drawn from a population-level distribution. The prior for all group parameters are assumed unit normal, with mean *μ* = 0 and standard deviation *σ* = 1. For learning rate, the parameter fitting was done in the untransformed space (−∞, +∞) and transformed to [0, 1] using the unit normal cumulative distribution function (CDF). We compute confidence (technically credible) intervals and statistical significance from percentiles of the posterior samples over the group-level parameters (or functions of them: specifically, their difference). In particular we compute a statistic comparable to a one-or two-tailed *P-*value as one minus (twice) the size of the largest credible interval that encompasses the point hypothesis (e.g. zero).

### Multivariate MEG Analysis: Stimuli Decoding and Sequences

Sequenceness analysis relies on an ability to quantify transient spontaneous neural reactivations of task stimuli. For each stimulus (*k* ∈ [1: 18]) indicating intermediate states in the RL task (e.g., *A*1, *A*2, *A*3, *B*1, *B*2, *B*3, etc), we trained a separate lasso-regularised logistic regression model based on their evoked neural response in the Functional Localizer task. Each model *k* discriminated between sensor patterns pertaining to stimulus *k* compared to all other stimuli plus an equivalent amount of ‘null’ data from the inter-trial interval. Inclusion of null data reduces the spatial correlation between classifiers, and helps sequence detection (*15*, *16*). To quantify classifier accuracy, the models were trained in leave-one-out cross-validation scheme and prediction accuracy estimated as the average proportion of test trials where the classifier reporting the highest probability corresponded to the trial label (**Fig 2A and** **Fig S1**). The cross-validation accuracy peaked around 200 ms, which is similar to our previous studies using a similar set of stimuli (*16*, *17*), and is also the time bin reported to give the strongest semantic representation (*38*). We confirmed that decoding accuracy was significantly greater than chance level using nonparametric permutation test. Specifically, we permuted the labels of test trials 500 times, and identified the maximal mean accuracy over time from 0 to 800 ms post-stimulus onset (controlling for multiple tests over time). Accuracy in the unpermuted data was considered significant if it exceeded 95% percentile on the maximal accuracy obtained from the permutation samples (dotted line in **Fig 2A and** **Fig S1**).

We trained the stimuli classifiers based on the whole brain multivariate sensor pattern at 200 ms post-stimulus onset (**Fig S1**). These classifiers will be used to decode state reactivations in the later RL task for sequence analysis. L1 regularisation, *λ*, was used to encourage sparsity and enhance sensitivity for sequence detection (*15*, *16*). To ensure the results were not overfit to the regularization parameter, we fixed *λ* = 0.005 for all subjects, based on sequence results on a pilot subject (which was not included in formal analysis). On the pilot data, this *λ* value maximizes an average sequenceness value across 10 ms to 200 ms state-to-state time lags.

We applied the 18 trained classifiers to each trial in the RL task, after outcome value receipt, this gave us a time*state decoding matrix trial by trial. We then used Temporally Delayed Linear Modelling (TDLM) to quantify evidence of sequential reactivations in this decoding matrix (*15*). This is the same analysis approach we have applied in previous studies, and is able to quantify sequenceness in forward and backward direction separately, while controlling for auto-correlation (*16*, *17*). We quantified sequenceness for all possible transitions permitted by the task structure. Evidence of replay in all 6 paths were estimated at the same time, and thereby controlled for common variance. We calculated sequenceness from time lag 10 ms to 500 ms.

The statistical significance was assessed using a state-identity based permutation test (*15*). The null hypothesis is that the state identities are exchangeable. In these permutations, we measure sequenceness of random transitions that are not consistent with the task structure, e.g., *A*1 → *B*3, also from 10 ms time lag to 500 ms. The permutations were run 1000 times. We determined the significance threshold by first taking the maximum sequenceness in the permutations across all computed time lags (to control for multiple comparisons), and then the 95% percentile on that peak across samples. Any true sequenceness that exceed this threshold is deemed significant. This approach has been validated in previous work in both simulation and empirical data (*15*–*17*).

### Sequence - Behavior Modelling

To test whether non-local replay facilitates non-local learning we again modified the baseline Q-learning model by sorting nonlocal paths into two categories and estimating separate learning rates: *α*_*replay*_ for paths with significant replay and *α*_*no-replay*_ for paths without significant replay. The learning rules are as before (except that both or neither non-local paths might have significant replay on a trial).

We identified significant replay events, in each path and at each trial. Significance is determined as before, except we now assess significance separately for each path and each trial, rather than as a grand average. We do this separately for 30 ms forward and 160ms backward sequences.

The other aspects of the analysis including the softmax decision rule, model prior and fitting procedure were exactly the same as described above. In this modelling analysis, we are interested in whether *α*_*replay*_ > *α*_*no-replay*_.

## Supplementary Text

### Sequenceness as a function of reward and starting arms

In addition to model-based analysis, we also asked how replay differs when responding to reward vs. no reward, and whether it is modulated by starting arm probability. We reasoned that if replay is indeed sensitive to the gain from policy improvement, rather than prediction error, all else being equal, replay should be stronger in trials where subjects received no reward, since those trials have a higher chance of necessitating a change of action in the next trial than trials where subjects received a reward. This reasoning is further supported by modelling results showing that the probability of having higher gain paths for non-local experiences is higher in trials when the subject does not receive a reward (*P*(*high gain*|*No reward*) = 0.73), compared to when receiving a reward (*P*(*high gain*|*reward*) = 0.32). We found that the 160 ms backward replay of non-local paths are also higher for no-reward trials compared to reward trials (**Fig S3**). This reward modulated replay is also stronger in high need (common) arm, compared to low need (rare) arm (**Fig S3**). Those results were specific to replay of nonlocal paths leading to the same end state, with no significant differences found for paths leading to a different end state (**Fig S3**). Although alternative explanations might exist, one possible account is that this reward modulated replay is prioritized based on need.

### Reactivation analysis

In addition to sequence analysis, we also examined the role of reactivation alone. In theory, subjects could reactivate only the first stimuli of paths during credit assignment time for updating the value, given that the main RL task contains requires only one action and, therefore, does not really require sequencing. Choice is made on the first state of paths under each arm. This type of reactivation account has been suggested on the basis of previous studies using sensory preconditioning paradigms over associations consisting of only two stimuli (*5*, *38*). We did not find evidence for this in our data. Reactivation of the first stimuli of paths did not facilitate learning, nor were they modulated by reward, choice, or arm (**Fig S4**). We have also looked at arm pictures during credit assignment time and again did not find reactivation of an arm facilitates learning, nor was it related to reward, or choice (**Fig S4**). Note we could effectively decode all three arms (with a peak cross-validation accuracy around 54±1.3%, **Fig S4**). We used the same training time (200 ms) and L1 regularization as we used throughout out the paper (results shown in Fig S4). We have also tested both L1 and L2 regularization as well as a wide range of regularization values for the reactivation analysis, but none give significant results. It is possible that different types of representation were reactivated and we have not tried training classifiers at different time bins, or in combinations with different regularization values (*38*). These null results indicate our sequential replay results cannot be explained by reactivation alone. It is also possible that although the RL task itself does not require sequencing, a prior extensive learning of task structure, and a requirement to always remember relational structure (i.e., sequences) throughout the experiment have encouraged subjects to choose a sequential mechanism, rather than reactivation alone.

### Reactivation and sequence analyses at decision time

In theory, this task can also be solved by prospective planning at decision time using either reactivation or sequential replay, i.e. at decision time, one can prospectively reactivate a desirable outcome state, or sequentially replay a path leading to this outcome. If either is the case, then reactivation or replay content in the decision time should predict what subjects are going to choose. This mechanism was suggested in a previous fMRI study (*4*). We did not find the reactivation outcome predicts behavior, nor was it modulated by reward (**Fig S5**), although we can decode outcome states well (with a peak cross-validation accuracy around 65±1.1%, **Fig S5**). We also found no evidence of sequential replay in general at decision time (**Fig S5**). This is in fact expected based on the model by Mattar and Daw (*20*), which suggests that both forward and backward replay can be used equivalently for learning and planning. Given that reward information can be fully incorporated at outcome time via backward replay, there is little cognitive imperative for prospective planning via forward replay.

---

**Fig. S1.**
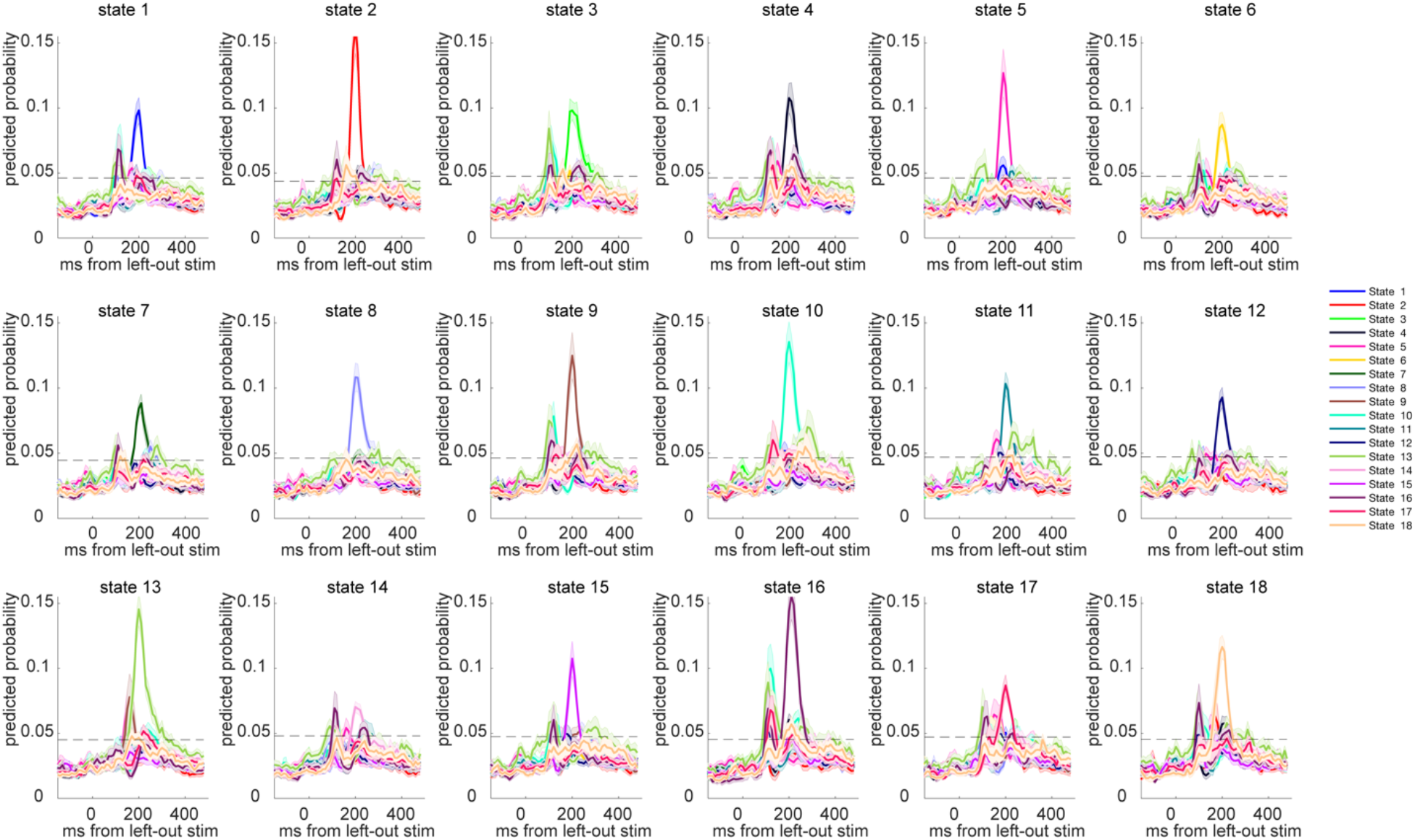
Classifier Performance of the Lasso Logistic Regression Models. Leave-one-out cross-validation results for each stimulus (18 in total), with classifiers trained and tested in functional localizer task. Dotted line indicates the permutation threshold estimated by randomly shuffling the labels and re-doing the decoding process. Classifiers were trained at 200 ms post stimulus onset, both in these plots as well as throughout the paper in reactivation and sequence analysis.

**Fig. S2.**
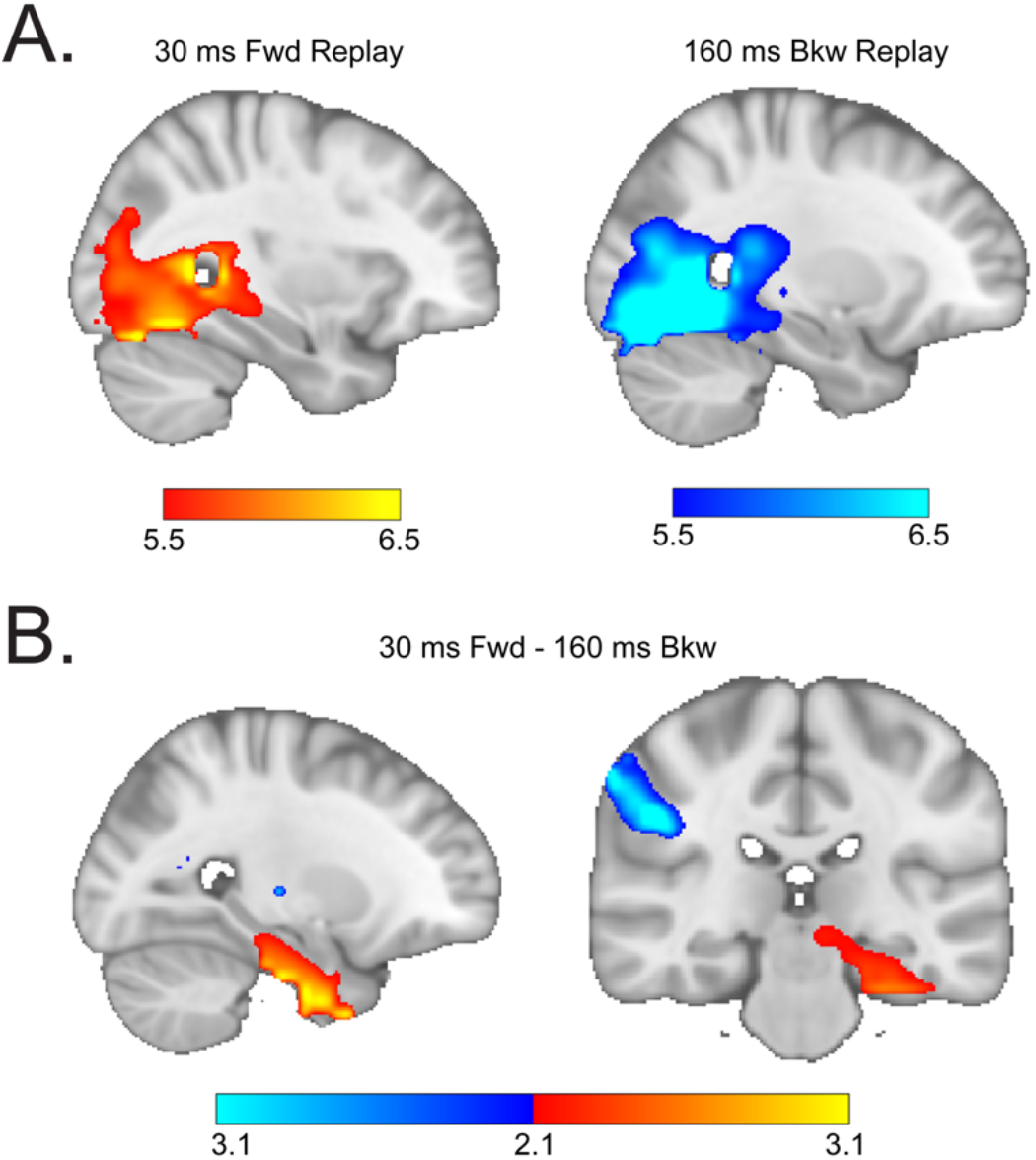
Whole-brain source localization of the two types of replay. **(A)** Source localization of the replay onset for the 30 ms forward replay and 160 ms backward replay separately, revealed significant visual and medial temporal lobe (MTL, including hippocampus) activation (peak Montreal Neurological Institute [MNI] coordinate: X = −17, Y = −45, Z = −16 for 30 ms forward replay, X = 18, Y = −66, Z = −16 for 160 ms backward replay). The activation in visual cortex and MTL encompass the same cluster and survived whole-brain multiple comparison correction based on a non-parametric permutation test (cluster forming threshold, *t* = 5, *n* =5000) for both 30 ms and 160 ms replay. **(B)** Contrast between 30 ms forward replay vs. 60 ms backward replay revealing higher activation in the MTL (peak MNI coordinate: X = −26, Y = −10, Z = −29) for 30 ms replay, and higher cortical regions −postcentral gyrus (peak MNI coordinate: X = 49, Y = −27, Z = 32) for 160 ms replay. Both the MTL (higher in 30 ms forward replay) and postcentral gyrus (higher in 160 ms backward replay) survived whole-brain multiple comparison correction based on a non-parametric permutation test (cluster forming threshold, *t* =2.1, *n* =5000).

**Fig. S3.**
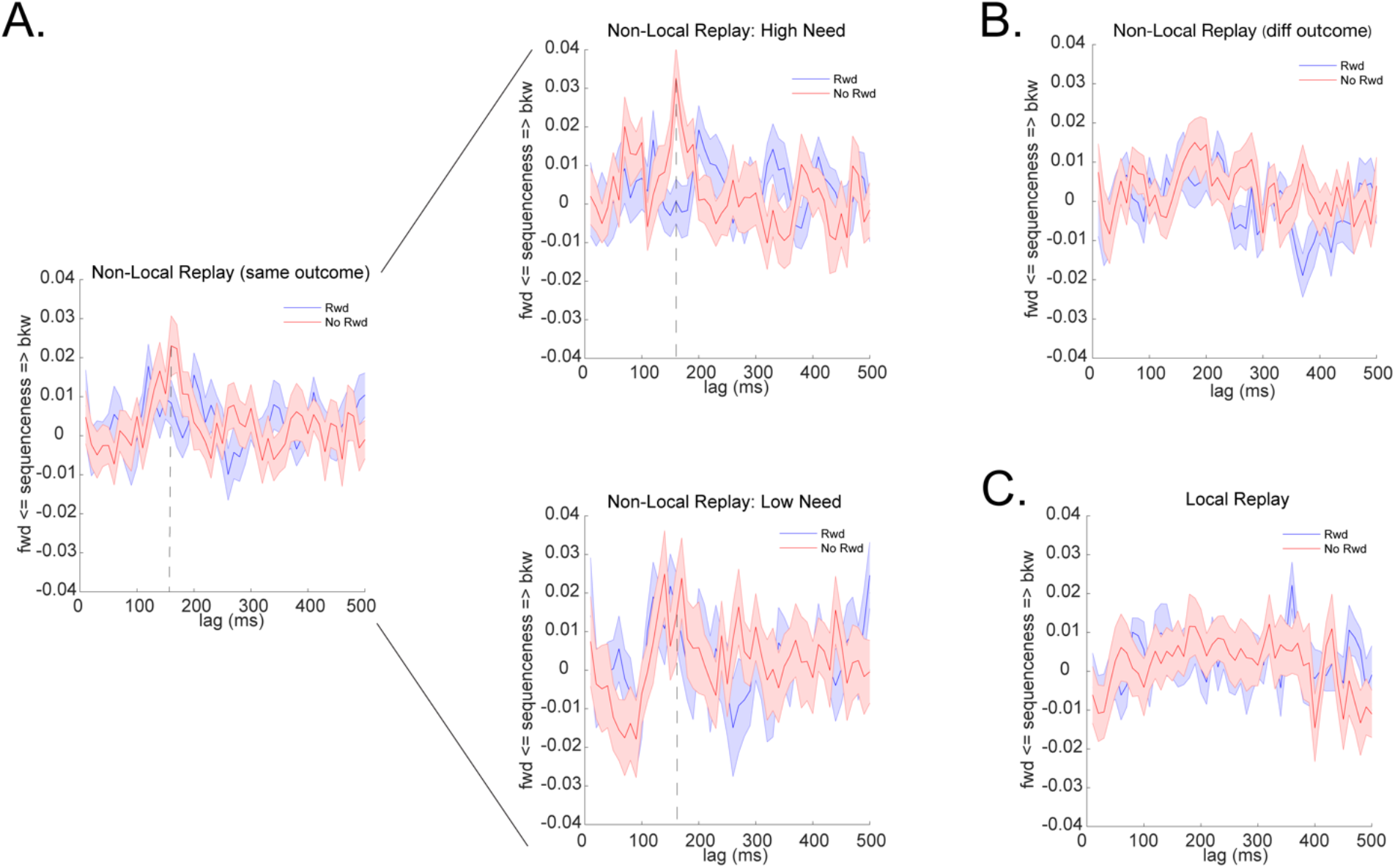
Non-local Replay is modulated by reward and need during credit assignment. **(A**) Stronger backward replay at 160 ms time lag in non-local paths that lead to the same end state, if this end state provided no reward vs. reward. This reward modulated replay is greater for the nonlocal path belonging to a high vs. low need arm. The x-axis shows time lag, and the y-axis shows the difference between backward (bkw) sequence and forward (fwd) sequence. Positive value indicates higher bkw than fwd replay, and vice versa. The dotted line indicates results at 160 ms time lag, which is the focus in the paper, and it is also the peak evidence for reward modulation. **(B)** This modulated replay does not exist for non-local path that leading to different outcomes, **(C)** Local replay is also not modulated by reward.

**Fig. S4.**
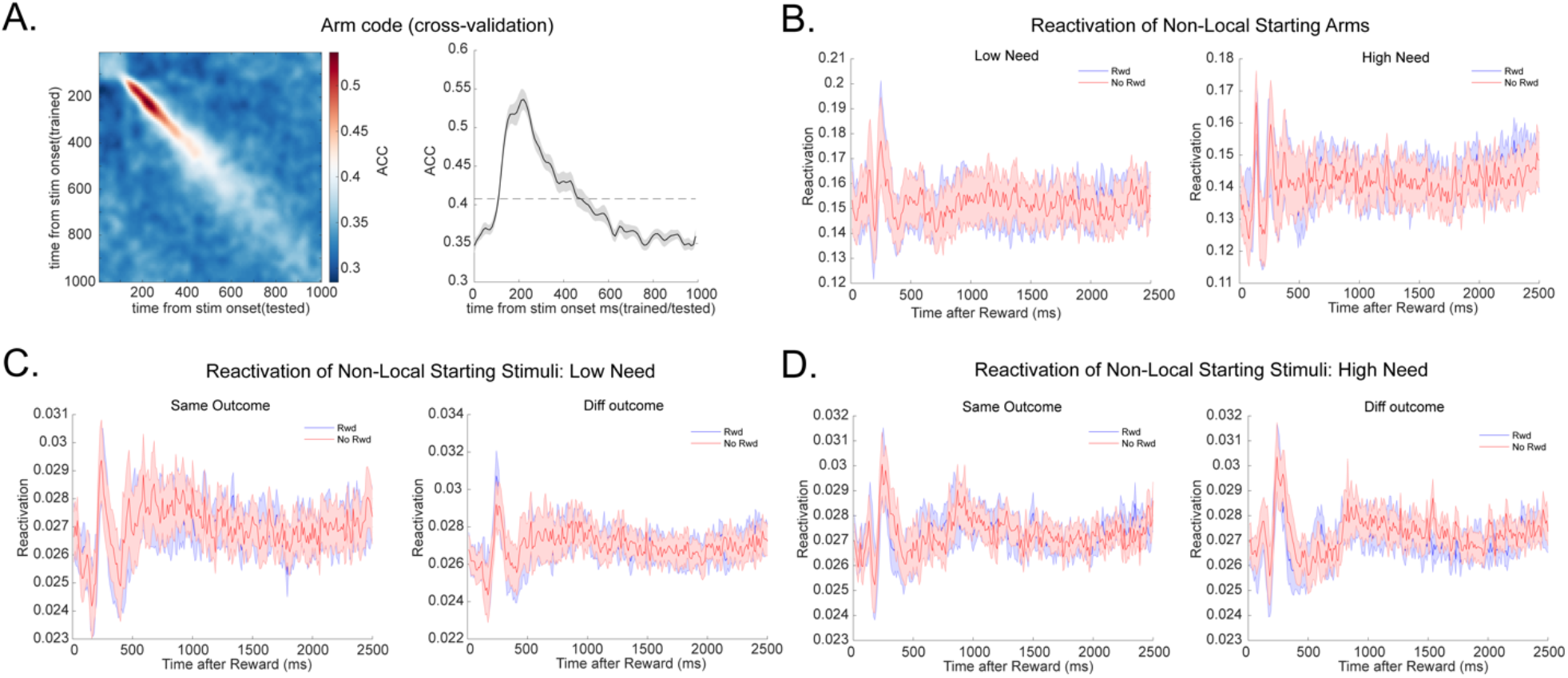
Reactivation of the starting stimuli and non-local arm during credit assignment. **(A)** Temporal generalization and decoding accuracy (leave one-out cross validation) for the three starting arms. We trained the arms classifiers based on the evoked neural response in the quiz of *arm learning* task where only one arm picture was presented in the centre of screen and the participants were required to think about which two paths belong to this arm. We observed a similar neural dynamic of the arm representation. We trained the arm classifiers in exactly the same way as the 18 stimuli classifiers. The dotted line is the permutation threshold. **(B)** Applying the trained arm classifiers to the outcome receipt time in the RL task, we see that reactivation alone of the non-local arms is not modulated by reward or need. (**C, D**) We have also looked at reactivation alone of the two starting stimuli which the participants are choosing from. They are also not modulated by reward, leading to same/different outcome, or need.

**Fig. S5.**
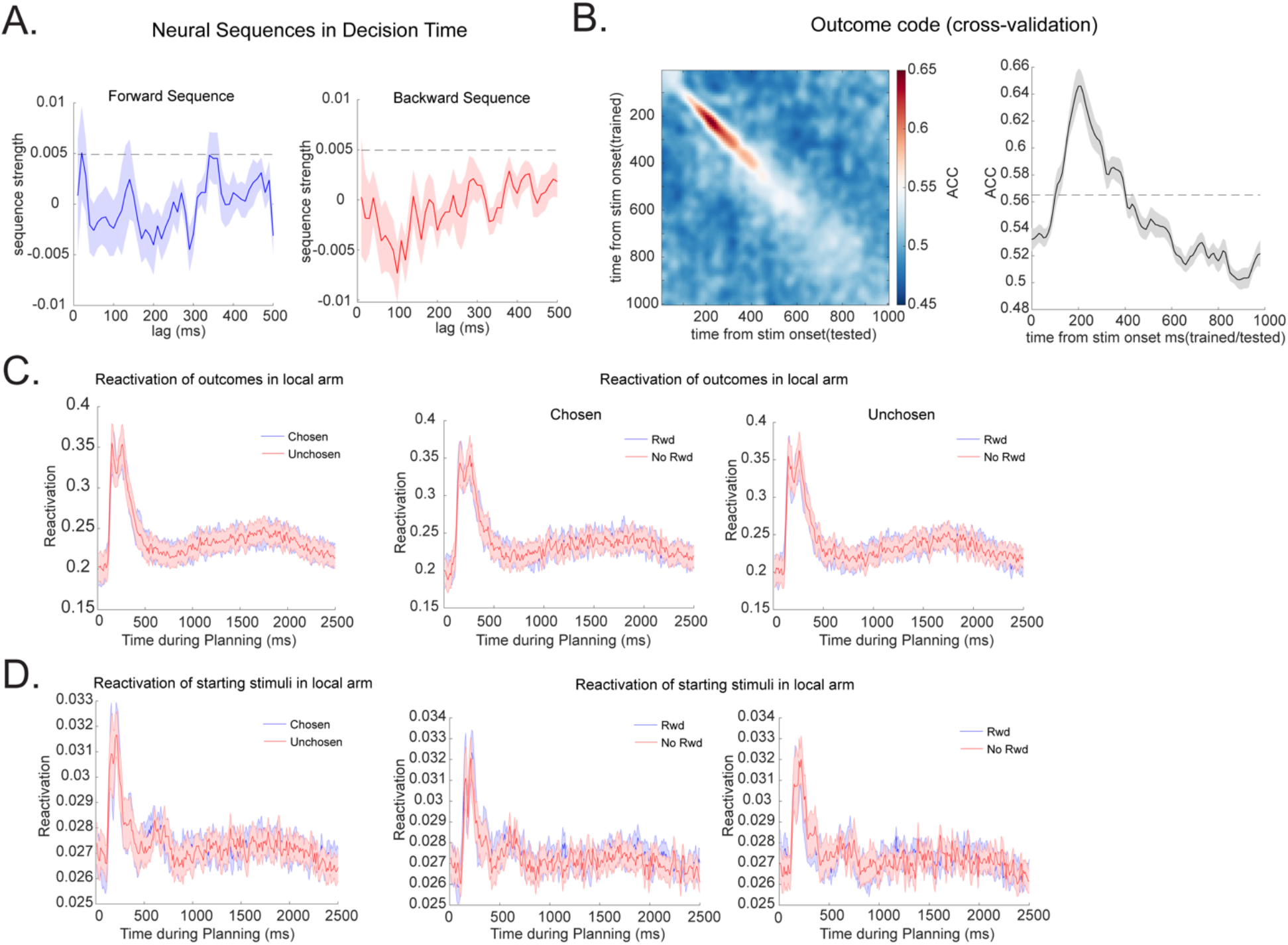
Replay and reactivation results at decision time. **(A)** No reliable neural sequence was seen at decision time. The only sequenceness scores that passed a permutation threshold is a 20 ms lag forward replay. This sequence is consistent with a faster replay (i.e., the 30 ms forward replay) we found during reward period, and with previous study (*16*). However, it is not modulated by either reward, need or choice. **(B)** Decoding results (leave one-out cross validation) for the two end states, which we call outcome code. We trained the outcome classifiers based on the evoked neural response at the quiz of the end state leaning task, where one end-state picture was presented in the centre of screen at a time, and subjects were asked to think about paths leading to this end. The dotted line is the permutation threshold. We again trained the outcome classifier in the same way as the arm and stimuli classifiers. **(C)** Applying the outcome classifiers to the decision time, we show that the end state reactivation is not modulated by reward or choice. **(D)** Reactivation alone of the two staring stimuli in the current arm was also not modulated by choice, reward, or need.

**Table S1.** **Estimates of free parameters (mean, 5% confidence interval and the potential scale reduction factor on split chains, 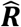) from the gain/need model.** Free parameters: ***α***_***d***_ – learning rate for direct experience, ***α***_***h***_ – learning rate for non-local experience of high gain (gain model) or need (need model), ***α***_***l***_ – learning rate for non-local experience of low gain (gain model) or need (need model), ***β*** - inverse temperature.

| Parameters | Gain model |  |  | Need model |  |  |
| --- | --- | --- | --- | --- | --- | --- |
| | Mean | 5% | $\hat{R}$ | Mean | 5% | $\hat{R}$ |
| $\alpha_d$ | 0.63 | 0.54 | 1.00 | 0.64 | 0.55 | 1.00 |
| $\alpha_h$ | 0.79 | 0.65 | 1.00 | 0.61 | 0.53 | 1.00 |
| $\alpha_l$ | 0.37 | 0.14 | 1.00 | 0.54 | 0.45 | 1.00 |
| $\beta$ | 1.40 | 1.20 | 1.00 | 1.40 | 1.10 | 1.00 |

**Table S2.** **Estimates of free parameters (mean, 5% confidence interval and the potential scale reduction factor on split chains, 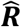) from the sequence (160 ms vs. 30 ms) - behavioral model.** Free parameters: ***α*_*d*_** – learning rate for direct experience (i.e., local path), ***α*_*replay*_** –learning rate for non-local path with replay, ***α*_*no-replay*_** – learning rate for non-local path without replay, ***β*** - inverse temperature.

| Parameters | 160ms backward replay |  |  | 30ms forward replay |  |  |
| --- | --- | --- | --- | --- | --- | --- |
| | Mean | 5% | $\hat{R}$ | Mean | 5% | $\hat{R}$ |
| $\alpha_d$ | 0.65 | 0.55 | 1.00 | 0.65 | 0.55 | 1.00 |
| $\alpha_{replay}$ | 0.70 | 0.56 | 1.00 | 0.61 | 0.50 | 1.00 |
| $\alpha_{no-replay}$ | 0.61 | 0.47 | 1.00 | 0.60 | 0.51 | 1.00 |
| $\beta$ | 1.40 | 1.20 | 1.00 | 1.40 | 1.10 | 1.00 |

## Notes

### Competing Interest Statement

The authors have declared no competing interest.

